# *MYC* overexpression leads to increased chromatin interactions at superenhancers and c-Myc binding sites

**DOI:** 10.1101/2021.01.04.425344

**Authors:** Yi Xiang See, Kaijing Chen, Melissa J. Fullwood

## Abstract

The MYC oncogene encodes for the c-Myc protein and is frequently dysregulated across multiple cancer cell types, making it an attractive target for cancer therapy. There have been many difficulties in targeting c-Myc, due to its complex network of regulators and the unstructured nature of its protein. Thus, we are interested in looking at the downstream cancer-specific functions of c-Myc. Overexpression of MYC leads to c-Myc binding at active enhancers, resulting in a global transcriptional amplification of active genes. However, the mechanism underlying this c-Myc enhancer invasion has not been well studied. To that end, we performed ChIP-seq, RNA-seq, 4C-seq and SIQHiC (Spike-in Quantitative Hi-C) on the U2OS osteosarcoma cell line with tetracycline-inducible MYC. MYC overexpression in U2OS cells modulated histone acetylation and increased c-Myc binding at superenhancers. SIQHiC analysis revealed increased global chromatin contact frequency, particularly at chromatin interactions connecting c-Myc binding sites. Our results suggest that c-Myc molecules are recruited to and accumulates within zones of high transcription activity, binding first at stable promoter binding sites at low expression levels, then at superenhancer binding sites when overexpressed. At the same time, the recruitment of c-Myc and other transcription factors may stabilize chromatin interactions to increase chromatin contact frequency. The accumulation of c-Myc at cancer-type specific superenhancers may then drive the expression of interacting oncogenes that each cancer is highly reliant on. By elucidating the chromatin landscape of c-Myc driven cancers, we can potentially target these chromatin interactions for cancer therapy, without affecting physiological c-Myc signaling.

## Introduction

Dysregulation of the *MYC* oncogene, which encodes for the c-Myc transcription factor, is common in cancer. Elevated mRNA and protein expression of *MYC* is seen across most cancer types in The Cancer Gene Atlas (TCGA) dataset (Schaub et al. 2018), making it an attractive target for cancer therapy. However, there are a multitude of causes for *MYC* dysregulation, including focal amplification of the *MYC* gene (Beroukhim et al. 2010), aberrations in the web of oncogenic signalling pathways regulating *MYC* (Kress et al. 2015), and the acquisition of new enhancer regulatory elements in spatial proximity to the *MYC* gene locus through translocation or overexpression of other transcription factors. The c-Myc protein itself is notoriously termed as “undruggable”, because of the unstructured nature of the protein and the lack of enzymatic binding sites or prominent pockets for small molecule inhibitor binding (McKeown and Bradner 2014).

c-Myc usually binds to the promoters of actively transcribing genes at physiological levels. Upon supraphysiological overexpression, c-Myc not only increases binding at active promoters, but “invades” into active enhancers as well, resulting in a global transcriptional amplification of active genes (Lin et al. 2012). Canonical c-Myc binding sites become saturated and c-Myc occupies non-canonical binding sites with lower affinity, resulting in an upregulation of genes associated with malignant transformation (Walz et al. 2014). Enhancer invasion has also been observed in the MYC family member N-Myc (Zeid et al. 2018).

At present, the mechanism underlying c-Myc enhancer invasion has not been well studied. Enhancers recruit transcription factors to activate linearly distant gene promoters via long-range chromatin interactions (Carter et al. 2002; Plank and Dean 2014). Recent studies have defined a category of enhancers known as superenhancers: clusters of enhancers with exceptionally high enrichment of transcriptional activators (Hnisz et al. 2013; Loven et al. 2013; Whyte et al. 2013). Superenhancers are associated with a more extensive network of chromatin interactions to multiple distant target genes (Cao et al. 2017), to serve as regulatory hubs to govern processes important to cell identity (Whyte et al. 2013). We asked whether c-Myc enhancer invasion might preferentially occur at superenhancers, and if this might lead to increased chromatin interactions.

In this study, we show that overexpressed c-Myc invades into superenhancers in particular. Using Spike-In Quantitative Hi-C (SIQHiC), we show that *MYC* overexpression leads to a global increase in chromatin contact frequency and increases significant chromatin loops at pre-existing c-Myc binding sites and superenhancers, suggesting that c-Myc superenhancer invasion may stabilize transient chromatin interactions to differentially regulate gene expression in cancer. Taken together, our results provide new insights into epigenomic consequences of c-MYC overexpression.

## Results

### *MYC* overexpression leads to increased c-Myc binding at superenhancers

We profiled the c-Myc enhancer binding landscape changes associated with *MYC* overexpression using the U2OS osteosarcoma cell line inserted with a tetracycline-inducible *MYC* cassette, because these cells are not addicted to *MYC* expression and have been used in previous studies to investigate the effects of *MYC* overexpression (Walz et al. 2014; Lorenzin et al. 2016). Doxycycline treatment for 30 hours increased *MYC* gene expression by about 35 times compared to vehicle treatment, together with an increase in c-Myc protein expression (Supplemental Fig. S1A-C). Hereafter, we refer to vehicle-treated and doxycycline-treated U2OS cells as Low *MYC* cells and High *MYC* cells respectively. RNA-seq identified 549 upregulated and 842 downregulated transcripts after *MYC* overexpression (Fig. 1A). 24.5% of the upregulated transcript promoters were bound by c-Myc at endogenous *MYC* expression levels, and increased to 36.9% after *MYC* overexpression (Fig. 1B). In contrast, only 12.7% of the downregulated transcripts were bound by c-Myc after overexpression, suggesting that the majority of the downregulated transcripts were secondary targets of c-Myc (Fig. 1B).

**Figure 1.**
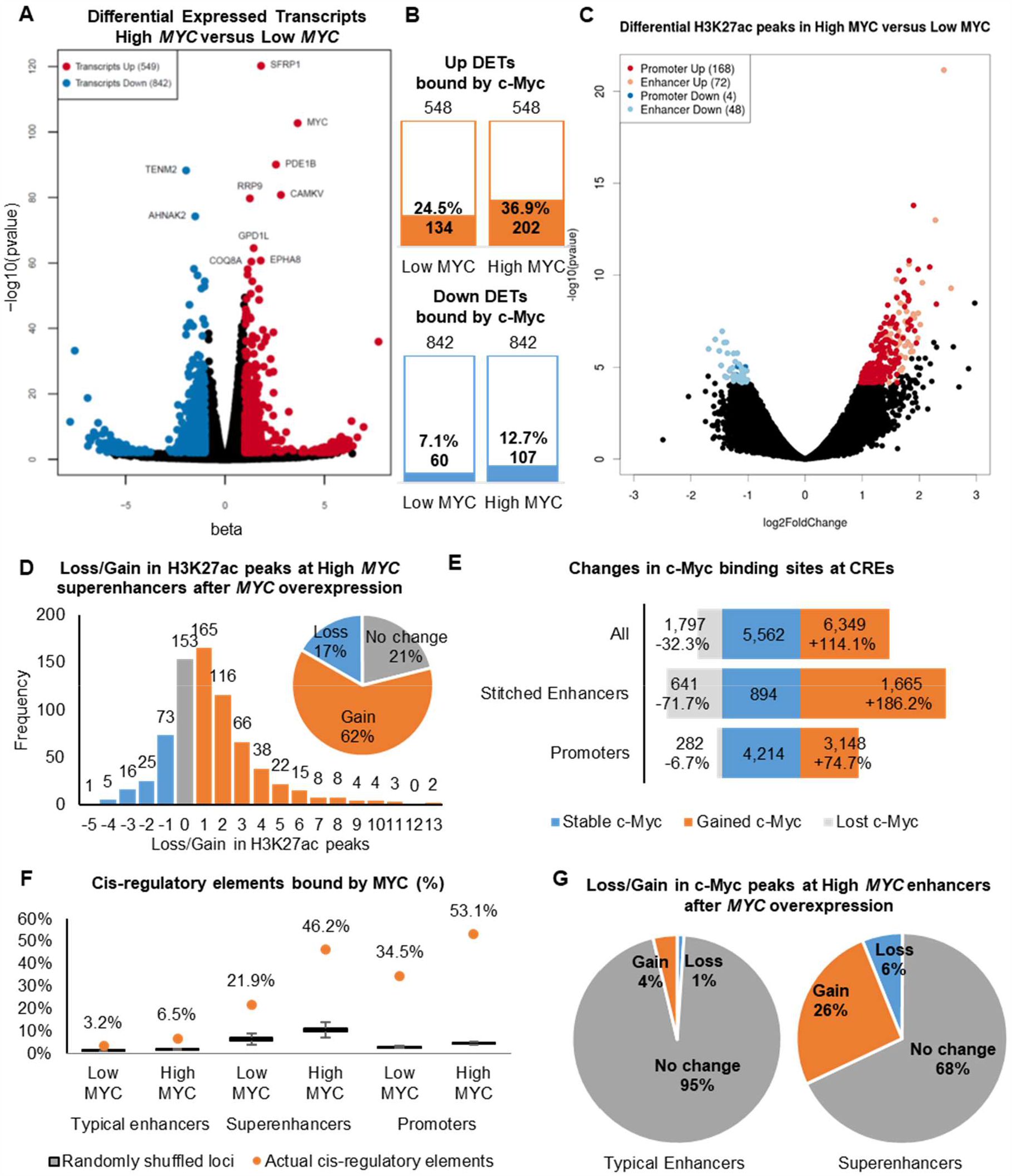
*MYC* overexpression leads to differential transcript expression, differential H3K27ac signal and increased c-Myc binding at superenhancers. (**A**) Volcano plot showing significantly upregulated (red) and downregulated (blue) transcripts after *MYC* overexpression (| beta| >1,FDR<0.01, Wald test). Top 10 significantly regulated transcripts are labelled. (**B**) Number of differentially expressed transcripts bound by c-Myc in Low *MYC* and High *MYC* cells. (**C**) Volcano plot showing H3K27ac peaks with significant changes in H3K27ac signal. (| log2foldchange| >1, FDR<0.1, Wald Test). (**D**) Bar graph and pie chart showing gain and loss of H3K27ac peaks at superenhancers after *MYC* overexpression. (**E**) Bar graph showing gain and loss of c-Myc binding sites at stitched enhancers and promoters after *MYC* overexpression.1 (**F**) Proportion of typical enhancers, superenhancers and promoters being bound by c-Myc compared to randomly shuffled coordinates. Proportion of c-Myc bound CREs are shown as orange dots. Box plots show 1000 iterations of c-Myc occupancy at random genomic loci of the same size and on the same chromosome as the actual CREs. (**G**) Pie charts showing gain and loss of c-Myc ChIP-seq peaks at typical enhancers (left) and superenhancers (right) after *MYC* overexpression.

We performed H3K27ac ChIP-seq to identify active cis-regulatory elements (CREs) in Low *MYC* and High *MYC* cells. H3K27ac peaks within 2.5 kb of transcription start sites were labelled as active promoter peaks, while the remaining peaks were labelled as enhancer peaks. *MYC* overexpression significantly regulated 292 H3K27ac peaks (168 increased and 4 decreased promoter peaks, 72 increased and 48 decreased enhancer peaks) (Fig. 1C), suggesting that *MYC* concentration modulates epigenetic regulation of transcription. We identified superenhancers using a similar approach to ROSE (rank ordering of superenhancers) (Loven et al. 2013; Whyte et al. 2013). Briefly, enhancer peaks within 12.5 kb of each other were stitched together, and superenhancers were separated from typical enhancers based on their H3K27ac ChIP-seq signal (Methods, Supplemental Fig. S1D). In total, we identified around 28,500 stitched enhancers in both Low *MYC* and High *MYC* cells, of which around 7,300 stitched enhancers were lost and gained after *MYC* overexpression (Supplemental Fig. S1F). We identified 890 superenhancers in the Low *MYC* cells and 725 superenhancers in the High *MYC* cells (Supplemental Fig. S1H). Almost all High *MYC* superenhancers overlapped with Low *MYC* stitched enhancers (Fig. 1D). 62% of the High *MYC* superenhancers gained between 1 and 13 H3K27ac peaks after *MYC* overexpression, with 15% merging multiple Low *MYC* stitched enhancers (Fig. 1D, Supplemental Fig. 1I), resulting in superenhancers with significantly more constituent H3K27ac peaks (Supplemental Fig. 1J).

Next, we looked into c-Myc occupancy at CREs. *MYC* overexpression led to a loss of 1,797 c-Myc binding sites and a gain of 6,349 c-Myc binding sites, of which 3,148 binding sites were gained at promoters and 1,665 at stitched enhancers (Fig. 1E). The proportion of c-Myc binding sites at stitched enhancers increased from 17.6% to 21.4%, whereas the proportion of c-Myc binding sites at promoters remained similar (61.5% to 61.8%) (Supplemental Fig. 2A), recapitulating previous observations of enhancer invasion. Strikingly, 21.9% of superenhancers overlap with c-Myc binding sites, increasing to 46.2% after *MYC* overexpression (Fig. 1F). 26% of superenhancers gained c-Myc binding sites after MYC overexpression, compared to only 4% of typical enhancers (Fig. 1G, Supplemental Fig. S2B-C). H3K27ac ChIP-seq profiles of c-Myc binding sites show that c-Myc binds directly at the constituent H3K27ac peaks and not randomly within the superenhancers (Supplemental Fig. S2D). To test whether the preferential binding of c-Myc at superenhancers was due to size, we randomly shuffled the genomic intervals of CREs within the same chromosome and overlapped them with c-Myc binding sites. Significantly more actual superenhancers overlapped with c-Myc binding sites than 1000 iterations of randomly shuffled genomic loci (Fig. 1F), indicating that c-Myc preferentially binds at superenhancers regardless of their size. Together, these results indicate that overexpressed MYC preferentially accumulates at H3K27ac peaks in superenhancers.

We further compared the c-Myc binding sites that were lost, gained or remained stable after *MYC* overexpression. Stable c-Myc binding sites at promoters and enhancers consistently have the highest H3K27ac and c-Myc ChIP-seq signals. Intriguingly, lost enhancer c-Myc binding sites had the lowest average H3K27ac peak signal (Supplemental Fig. S2E-F), but lost promoter c-Myc binding sites had high H3K27ac signal compared to stable and gained c-Myc binding sites (Supplemental Fig. S2G). This suggests that supraphysiological c-Myc binding at enhancers is dependent on open chromatin (demarcated by H3K27ac), whereas supraphysiological c-Myc binding at promoters is dependent on other factors, such as binding site affinity and recruitment by other transcription factors (Lorenzin et al. 2016).

### Spike-in Quantitative Hi-C (SIQHiC) reveals increased global chromatin contact frequency per cell after *MYC* overexpression

In traditional Hi-C techniques, cross-sample normalization is based on the assumption that chromatin interactions are largely stable across biological conditions (Lun and Smyth 2015; Stansfield et al. 2018). However, perturbation of factors involved in chromatin organization such as CTCF and cohesin can result in global changes in chromatin contact frequency. We hypothesized that increased global c-Myc binding at CREs not only increases transcription of associated genes but also stabilizes chromatin interactions at these genes, resulting in similar global changes in chromatin contact frequency.

Cell-count normalization techniques have been developed for RNA-Seq (Lovén et al. 2012) and ChIP-Seq (Orlando et al. 2014) to account for global changes in transcription and occupancy respectively. Cell-count normalized transcription analyses had previously revealed a global increase in transcription after *MYC* overexpression (Loven et al. 2012; Nie et al. 2012). These methods involve mixing a control sample from an orthogonal species into the experimental human samples at a fixed cell number ratio. The recently published AQuA-HiChIP protocol (Absolute Quantification of Architecture Hi-ChIP) (Gryder et al. 2019) utilizes the same strategy of spiking in mouse cells into human samples, running on the assumption that contact frequency for untreated mouse cells should be the same across samples.

In our study, we adapted the AQuA-HiChIP protocol to perform Spike-In Quantitative Hi-C (SIQHiC) on Low *MYC* and High *MYC* cells in duplicate, to elucidate the global changes in chromatin contact frequency. In brief, mouse 3T3 cells were cross-linked and mixed into each sample of human cells at a ratio of 1:4, before continuing with the Arima Hi-C kit protocol (Fig. 2A, Methods). If global chromatin contact frequency is unchanged, we expect to obtain a similar human to mouse chromatin contact ratio (HMR) across all samples. Hence, the relative difference between the HMR of different samples reflects the differences in global chromatin contact frequency and can be used to normalize the samples (Fig. 2B).

**Figure 2.**
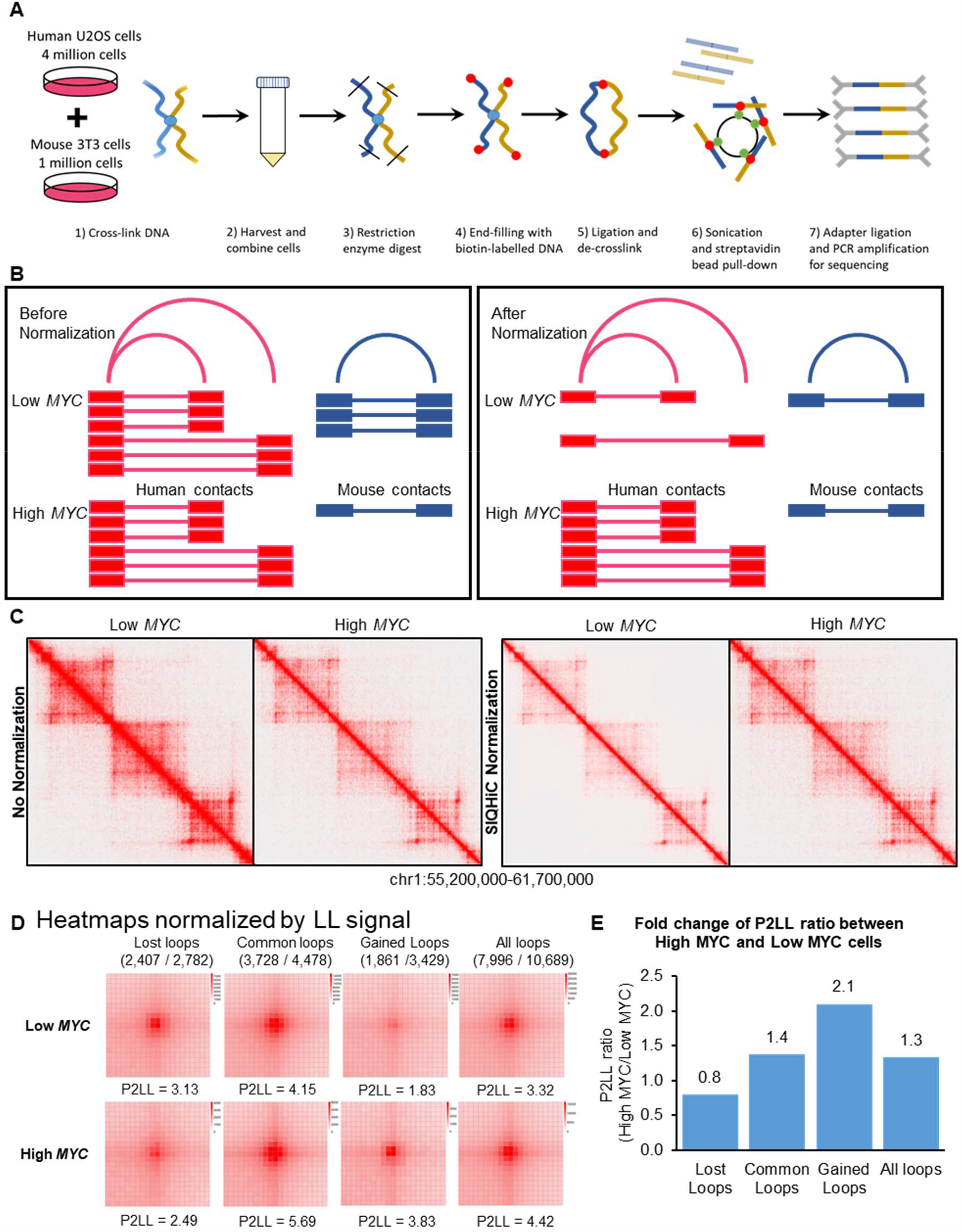
SIQHiC (Spike-In Quantitative Hi-C) normalization reveals increased chromatin contact frequency per cell after *MYC* overexpression. (**A**) Brief overview of the SIQHiC workflow. (**B**) Cartoon illustrating Hi-C contacts before (left) and after (right) SIQHiC normalization. SIQHiC normalization scaled down the Low *MYC* contacts such that the number of mouse contacts in both conditions were the same, thereby revealing an increase in human chromatin contacts. (**C**) Hi-C matrix heatmaps of a region on chromosome 1. Left panel: No normalization. Right panel: SIQHiC normalization. (**D**) Aggregate Peak Analysis (APA) plots at 5kb resolution showing the aggregate signal of “Lost”, “Common” and “Gained” chromatin loop sets in Low MYC and High MYC cells identified using the non-normalized Hi-C contact matrices. P: Peak signal at the centre pixel. LL: Average signal of the 3 x 3 square at the lower left corner of the APA plot, representing local background. P2LL: Ratio of P to LL. APA colour scales were normalized by the LL signal. Loop sets were filtered to remove short loops near the diagonal (shown above each APA plot; numerator: number of filtered loops; denominator: total number of loops). (**E**) Bar graph showing fold change of P2LL ratio between High *MYC* and Low *MYC* cells.

Ambiguous read pairs mapping to both species consistently made up a small percentage of the total reads per sample (∼0.3%) and were discarded (Supplemental Table S2). Using SIQHiC, we observe a 2.1 to 4.4 fold increase in HMR after *MYC* overexpression, suggesting an equivalent increase in global chromatin contacts (Supplemental Table S2). Hi-C chromatin interaction heatmaps of Low *MYC* and High *MYC* cells appear to show a decrease in chromatin interactions after *MYC* overexpression (Fig. 2C, left). After SIQHiC normalization, we observed an increase in chromatin interactions instead (Fig. 2D, right). As expected, Low *MYC* duplicates had similar HMR (33.4 and 36.4), reflecting consistent chromatin contact frequencies at physiological *MYC* levels (Supplemental Table S2). Conversely, High *MYC* duplicates had higher and more variable HMR (146.9 and 78.1) after doxycycline induction of *MYC* expression. (Supplemental Table S2).

### *MYC* overexpression does not affect compartments and TADs but strengthens a set of chromatin loops

Given that *MYC* overexpression increased chromatin contact frequency, we wanted to know whether *MYC* overexpression reshapes the global chromatin landscape. A/B compartment analysis showed few instances of compartment switching between Low MYC and High MYC cells, with chromosome 8 shown as an example in Supplemental Fig. S3A. We also identified similar numbers of topologically associating domains (TADs) between Low *MYC* and High *MYC* cells at 50kb (1,951 and 1,965) and 100kb resolutions (975 and 836) (Supplemental Fig. S3B-C), with more than 60% overlap. These results indicate that *MYC* overexpression does not alter chromatin compartments and TADs.

Next, we identified 7,266 and 7,910 significant chromatin loops in Low *MYC* and High *MYC* cells respectively, with approximately 60% of these loops common between both conditions (Supplemental Fig. S3D). We performed an Aggregate Plot Analysis (APA) to show the aggregate signal of the chromatin loops that were “gained” and “lost” after MYC overexpression, or “common” between both conditions. In this analysis, the 105kb x 105kb contact matrices surrounding the midpoints of all loops in each loop set are overlapped such that the center pixel (P) of the APA plot shows the aggregate signal of the midpoints of these loops. The lower left corner of the plot (LL) represents the contact frequency for shorter interactions in the vicinity of the loop set and gives an indication of the local random interaction frequencies. The P2LL score (ratio of signal at P to the average signal at LL) can then be used to show the chromatin interaction frequency relative to the local background. Without SIQHiC normalization, APA plots show decreased peak signal enrichment (P) at chromatin loops after *MYC* overexpression (Supplemental Fig. S3E). Using SIQHiC to normalize for cell count revealed an 2.2 and 3.3 fold increase in peak signal enrichment (P) at common and gained loop sets instead (Supplemental Fig. S3F). Using P2LL scores to normalize for local interactions, we observe a similar increase in chromatin contact frequency at common and gained loop sets (1.4 fold and 2.1 fold respectively) (Fig. 2D-E). Hence, APA analysis showed that, after *MYC* overexpression, the increase in chromatin interaction frequency is not merely a general increase in chromatin movement, but focused at the set of gained loops.

We zoomed in on the *MYC* gene locus since *MYC* is known to be regulated by superenhancers through chromatin interactions (Shi et al. 2013; Herranz et al. 2014; Zhang et al. 2016; Bahr et al. 2018). We generated virtual 4C plots from our Hi-C data using the *MYC* gene promoter as the viewpoint, to visualize the chromatin interactions at this locus. Without normalization, virtual 4C showed a general decrease in chromatin interaction frequencies (Fig. 3A). However, SIQHiC normalization of the Hi-C reads revealed a general increase in virtual 4C signal instead (Fig. 3A).

**Figure 3.**
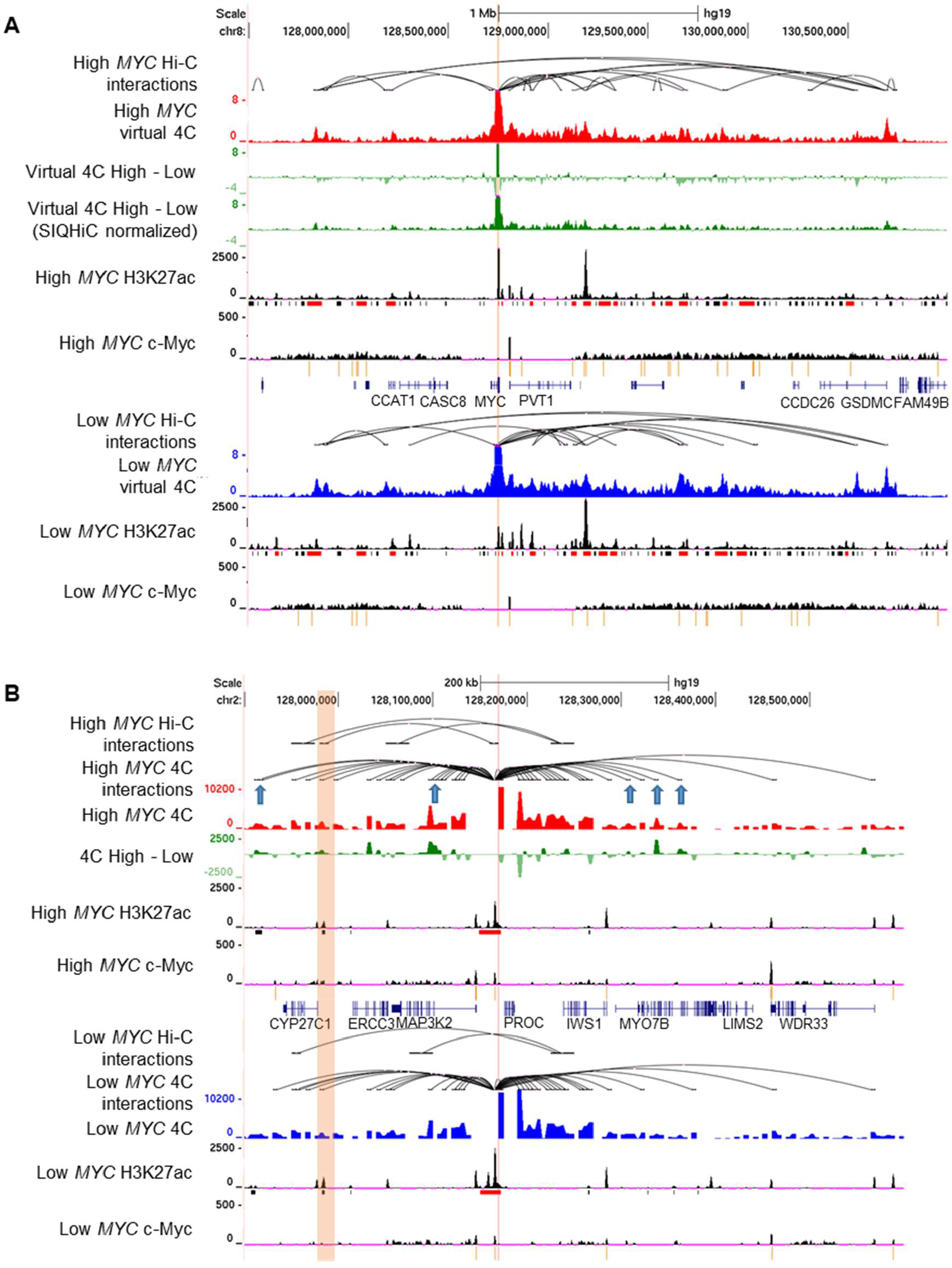
*MYC* overexpression increases chromatin contact frequency and chromatin loops at superenhancers. (**A**) Genome browser view of High *MYC* and Low *MYC* cells at the *MYC* locus Tracks show Hi-C chromatin loops, virtual 4C signal of interactions at the *MYC* promoter (red and blue tracks respectively), H3K27ac and c-Myc ChIP-seq signal. The difference between the virtual 4C signals of High *MYC* and Low *MYC* cells are shown with and without SIQHiC normalization (green tracks). Superenhancers (red) and typical enhancers (black) are shown as bars below the H3K27ac ChIP-seq tracks. c-Myc binding peaks are shown as yellow bars below the c-Myc ChIP-seq tracks. (**B**) 4C-seq of chromatin interactions at a randomly selected c-Myc bound superenhancer near the *PROC* gene (highlighted in pink) on chromosome 2. Tracks show Hi-C chromatin loops, 4C-seq signal (red and blue tracks), difference between 4C-seq signal of High *MYC* and Low *MYC* cells (green), H3K27ac and c-Myc ChIP-seq signal. Gained Hi-C chromatin loop is highlighted in orange. Blue arrows show additional gained chromatin interactions identified using 4C-seq.

Since Hi-C assays the entire genome, it requires very high sequencing depth for sufficient resolution and limits chromatin loop detection within a specific region of interest (Babu and Fullwood 2015; Sati and Cavalli 2017). Hence, we performed 4C-seq at a randomly selected c-Myc bound superenhancer near the *PROC* gene which gained a chromatin loop to the *CYP27C1* gene after *MYC* overexpression. 4C-seq showed that the gained Hi-C chromatin loop was already present in Low *MYC* cells, but *MYC* overexpression increased the interaction frequency of this loop (Fig. 3B). 4C-seq also identified additional chromatin loops at the c-Myc bound superenhancer locus that were not significant in the Hi-C data (Fig. 3B, blue arrows). Taken together, *MYC* overexpression strengthens a set of gained chromatin loops.

### Chromatin loops tend to connect c-Myc binding sites at superenhancers

We overlapped all Hi-C loop anchors with c-Myc binding sites and found that the proportion of all chromatin loops with c-Myc binding sites increased from 21.8% to 32.5% after *MYC* overexpression (Fig. 4A). Gained loops were particularly enriched in c-Myc binding, with 47.6% of these loops occupied by c-Myc in High *MYC* cells, which was considerably higher than the proportion of randomly shuffled chromatin loop coordinates overlapping with c-Myc binding sites (Fig. 4A). Notably, more gained loops were occupied by c-Myc at physiological c-Myc levels (34.3%) compared to common (16.8%) and lost loops (14.3%) (Fig. 4A). Additionally, 7.6% of the gained loops had c-Myc binding at both anchors at physiological c-Myc levels, and increased to 15.7% after *MYC* overexpression (Fig. 4B). These results suggest that chromatin interactions connecting c-Myc bound genomic loci are preferentially strengthened after *MYC* overexpression.

**Figure 4.**
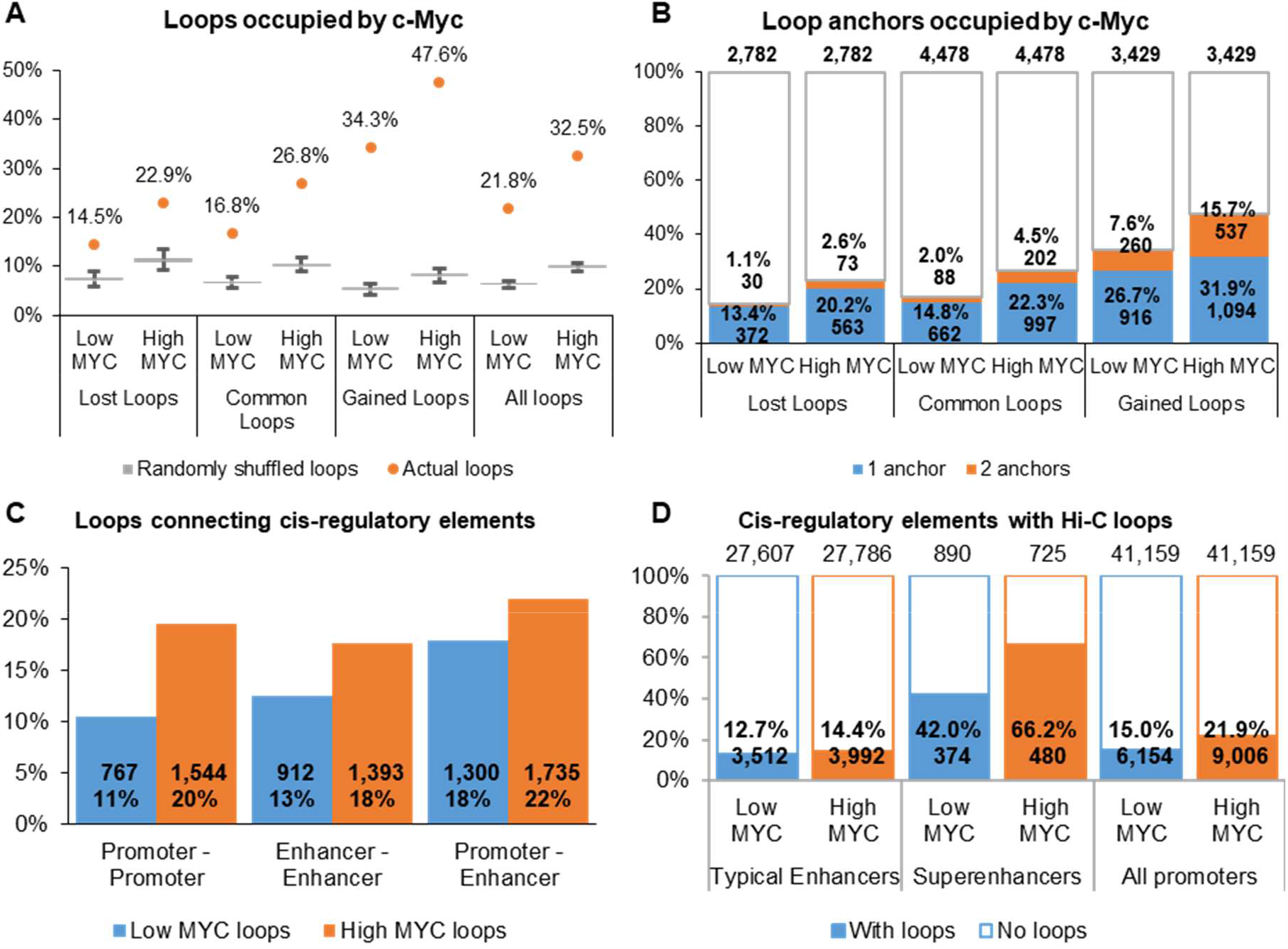
Chromatin interactions are enriched at superenhancers and gained chromatin loops tend to connect c-Myc binding sites. (**A**) Percentage of Hi-C chromatin loops occupied by c-Myc compared to randomly shuffled loop coordinates. Percentage of c-Myc bound chromatin loops are shown as orange dots. Box plots show 1000 iterations of c-Myc occupancy at random genomic loci of the same size and on the same chromosome as the actual chromatin loops. (**B**) Percentage of chromatin loop anchors occupied by c-Myc. (**C**) Percentage of Hi-C chromatin loops connecting cis-regulatory elements. (**D**) Percentage of cis-regulatory elements with Hi-C chromatin loops.

Next, we wanted to know whether chromatin interactions are specifically altered at the enhancers invaded by c-Myc. We annotated the Hi-C loops according to their overlap with promoters and stitched enhancers. *MYC* overexpression increased chromatin looping between promoters (11% to 20%), between stitched enhancers (13% to 18%) and between promoters and stitched enhancers (18% to 22%) (Fig. 4C). In particular, the proportion of CRE loops within the lost loop set was similar to the common loop set, whereas the proportion of CRE loops within the gained loop set was markedly higher (Supplemental Fig. S4A), showing that chromatin interactions are preferentially strengthened between CREs. Consistent with previous research (Cao et al. 2017), a markedly higher proportion of superenhancers (42.0 %) was associated with Hi-C loops compared to typical enhancers (12.7%) and promoters (15.0%), and *MYC* overexpression further increased the proportion of superenhancers with loops to 66.2% (Fig. 4D). c-Myc bound CREs were more associated with chromatin loops compared to non c-Myc bound CREs, particularly after *MYC* overexpression (Supplemental Fig. S4B). Seeing as superenhancers are larger than typical enhancers, we compared the constituent H3K27ac peaks within typical and superenhancers. Similarly, more c-Myc bound superenhancer constituent H3K27ac peaks overlapped with Hi-C loops (27.3% - 39.9%) compared to c-Myc bound typical enhancer constituents (13.2% - 21.2%) and promoter loci (15.4% - 28.0%) (Supplemental Fig. S4B-C).

Since c-Myc binds at both promoters and enhancers, we looked at promoter-enhancer chromatin loops (P-E loops) in detail. We overlapped P-E loops with c-Myc binding sites and categorized the loops as being bound proximal to the promoter loop anchor or bound at both loop anchors (Fig. 5A). *MYC* overexpression increased the proportion of P-E loops with only proximal c-Myc binding sites from 16% to 26%, while P-E loops bound at both anchors increased from 3% to 16% (Supplemental Fig. 4D). Although lost P-E loops gained c-Myc binding sites at the promoter proximal loop anchor after *MYC* overexpression (22% to 36%), few of these loci gained c-Myc at the distal anchor as well (3% to 8%) (Fig. 5B). On the other hand, gained P-E loops gained promoter proximal c-Myc binding sites (24% to 44%), but a larger proportion also gained distal c-Myc binding sites (9% to 23%) (Fig. 5B). Taken together, our results suggest that *MYC* overexpression preferentially strengthens chromatin interactions between c-Myc binding sites.

**Figure 5.**
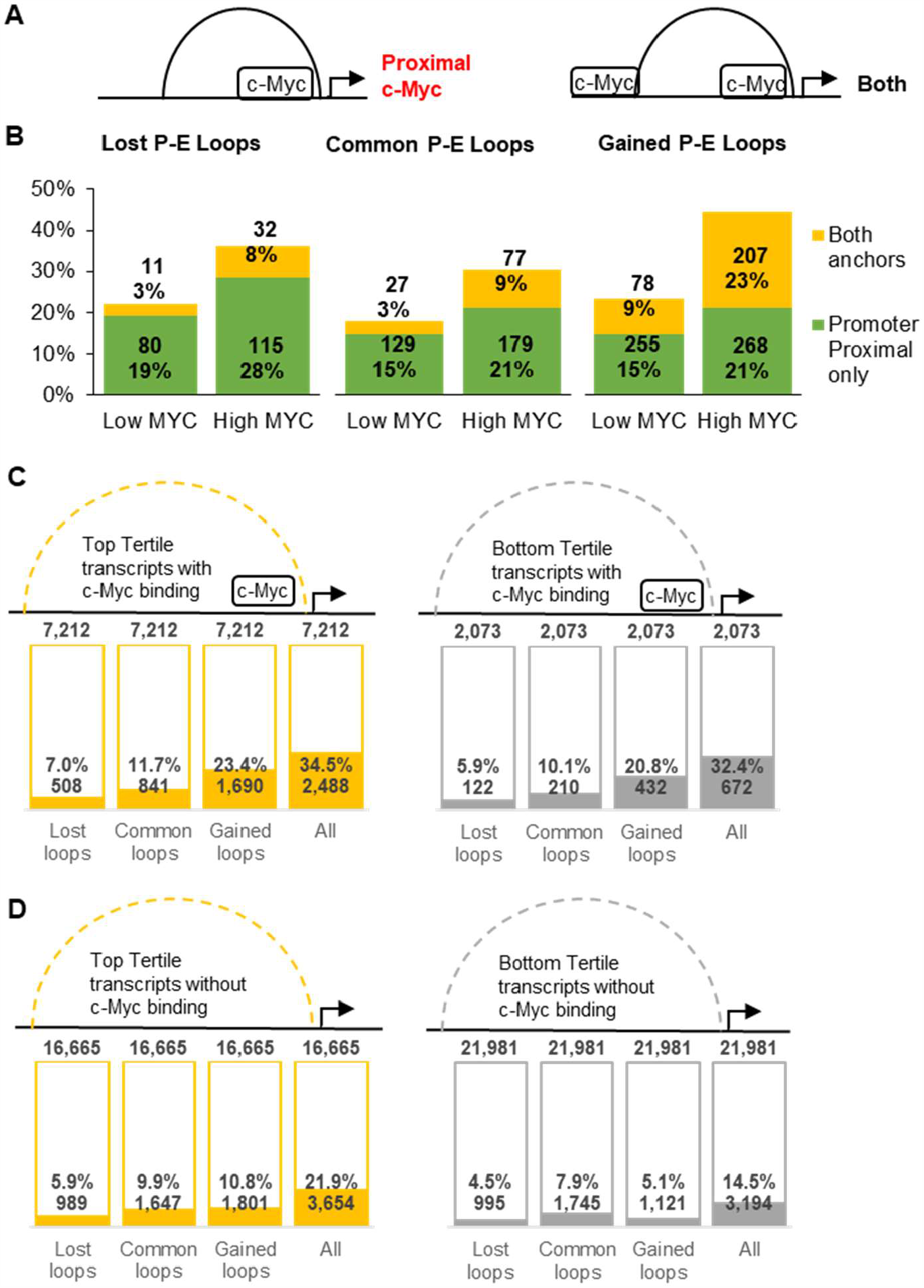
c-Myc accumulates at both anchors of gained promoter-enhancer chromatin loops. (**A**) Promoter-enhancer chromatin loops are categorized as having promoter-proximal c-Myc binding or c-Myc binding at both loop anchors. (**B**) Percentage of lost (left), common (middle) and gained (right) promoter-enhancer chromatin loops occupied by c-Myc at the promoter proximal anchor or at both anchors. (**C-D**) Percentage of (**C**) c-Myc bound and (**D**) non c-Myc bound transcripts with Hi-C chromatin loops. Transcripts were stratified into top (yellow), middle (not shown) and bottom (grey) tertiles based on transcript expression after *MYC* overexpression. (**E**) Proposed model of how *MYC* overexpression strengthens chromatin interactions at superenhancers. Superenhancers interact dynamically with promoters within a zone of high transcription activity. At physiological levels of c-Myc (left), c-Myc molecules occupies stable binding sites at promoters. *MYC* overexpression (right) leads to the accumulation of c-Myc molecules, occupying lower affinity binding sites at superenhancers and increasing the protein interactions within the zone, leading to reduced chromatin mobility and increased interaction frequency.

### Transcription activity does not affect chromatin interactions

Since c-Myc overexpression leads to a global increase in transcription, we wanted to know whether chromatin interactions are affected by these transcriptional changes. Firstly, we compared significantly up- and downregulated transcripts (P<0.05, abs(beta) >1) with non-regulated transcripts (P>0.95, transcripts per million>1). Hi-C loops are not preferentially gained or lost at *MYC* regulated transcripts compared to *MYC* non-regulated transcripts (Supplemental Fig. 5A). However, more c-Myc bound transcripts gained Hi-C loops (Supplemental Fig. 5B) compared to transcripts without c-Myc binding (Supplemental Fig, 5C), regardless of their differential transcript expression.

When transcripts were stratified into tertiles based on transcript expression, top tertile transcript promoters appeared to overlap better with Hi-C loops compared to the bottom tertile (25.7% and 16.1% respectively) (Supplemental Fig. 6A). However, this was primarily due to top tertile promoters being more enriched in c-Myc binding. c-Myc bound bottom tertile transcripts showed similarly high overlap with gained Hi-C loops as the c-Myc bound top tertile transcripts (20.8% and 23.4% respectively) (Fig. 5E), in contrast to bottom and top tertile transcripts without c-Myc (5.1% and 10.8% respectively) (Fig. 5F). These results show that it is the presence of c-Myc binding that increases chromatin interaction frequency and not transcription activity.

## Discussion

*MYC* overexpression has long been described as a key driving force for oncogenesis in multiple cell types, resulting in tumours that become highly dependent on elevated *MYC* expression(Gabay et al. 2014; Bradner et al. 2017), but it is still unclear how overexpressed c-Myc shifts from regulating its canonical target genes to activate oncogenes. Here, we show that *MYC* overexpression leads to superenhancer invasion and changes in the enhancer-ome and chromatin interactome. Chromatin interactions are strengthened at c-Myc binding sites, particularly at superenhancers, albeit with no significant changes in TADs or compartments.

Recent studies have shown that overexpressed c-Myc invades into distal enhancer regulatory elements to differentially regulate gene expression (Lin et al. 2012; Nie et al. 2012; Sabo et al. 2014). In this work, we show that *MYC* overexpression modulates histone acetylation to create superenhancers containing more constituent H3K27ac peaks and exquisitely high enrichment of c-Myc binding. Cancer cells have been shown to gain cancer-specific superenhancers that drive oncogene expression (Ooi et al. 2016; Tsang et al. 2019; Raisner et al. 2020). Here, although there was a loss and gain of more than 7000 stitched enhancers after *MYC* overexpression, we observed no *de novo* superenhancer formation. These results indicate that c-Myc invades into active superenhancers but is unable to activate silent oncogenic superenhancers by itself.

Since superenhancers are associated with abundant long-range chromatin interactions and high interaction frequencies (Schmitt et al. 2016; Beagrie et al. 2017; Cao et al. 2017), we wanted to know whether c-Myc superenhancer invasion alters the three-dimensional chromatin interactome. Using SIQHiC - a modified Hi-C protocol normalizing for cell count between samples, we observed increased global chromatin contact frequency after *MYC* overexpression. Chromatin interactions at c-Myc binding sites were preferentially strengthened and connected c-Myc binding sites together, notably between promoters and enhancers. Kieffer-Kwon et al. (2017) previously showed that *MYC* upregulation during B cell activation is accompanied by a two fold increase in chromatin loops, particularly the formation of B cell associated chromatin loops. In this work, *MYC* overexpression alone strengthened pre-existing loops but did not increase the total number of chromatin loops, suggesting that other chromatin factors are required to reconfigure the chromatin interaction landscape in cancer as well as in B cell activation.

Previous research has shown that active transcription may play a role in maintaining the three-dimensional genome organization (Busslinger et al. 2017; Isoda et al. 2017; Gu et al. 2018; Heinz et al. 2018). Here, we show that *MYC* overexpression increases chromatin contact frequency not because of increased transcriptional activity, but due to direct binding of c-Myc to the interacting loci.

c-Myc/Max heterodimers have been shown to form heterotetramers *in vitro* at physiological *MYC* conditions (Nair and Burley 2003). Additionally, phase separated condensates form at superenhancers to form high localized transcription factor concentrations (Sabari et al. 2018), and c-Myc molecules have been shown to participate in such phase-separated condensates (Boija et al. 2018). We speculate that c-Myc molecules are recruited to and accumulates within superenhancer phase-separated condensates with stable promoter c-Myc binding sites. Supraphysiological levels of c-Myc saturates the promoter binding sites and may then spill over to lower affinity binding sites at superenhancers. At the same time, promoter-superenhancer chromatin interactions with high c-Myc occupancy are preferentially strengthened by the interactions between multiple c-Myc molecules and other transcription factors such as the Mediator complex.

*MYC* overexpression is a common event in oncogenesis across multiple cancers, but cannot alone induce malignant transformation. Our results suggest that other cancer initiating mutations are required to reconfigure the epigenetic and chromatin interaction landscape, with c-Myc playing a crucial role in maintaining these epigenetic alterations by stabilizing these transient changes in chromatin interactions. As such, *MYC* addicted cancers may be exquisitely sensitive to inhibitors of architectural proteins such as CTCF and YY1. In particular, YY1 has been shown to mediate enhancer-promoter interactions (Weintraub et al. 2017) and binds in coordination with c-Myc (Shrivastava et al. 1996; Vella et al. 2012), making it a potential target for *MYC* addicted cancers.

Taken together, our manuscript has demonstrated that *MYC* overexpression leads to c-Myc invasion into superenhancers and SIQHiC uncovered an increase in chromatin interactions at these regions. SIQHiC will be useful for uncovering changes in chromatin contact frequency when perturbing other chromatin factors, such as architectural proteins and chromatin remodellers. Our results lay the groundwork for more research to be done to further elucidate the role of c-Myc in maintaining cancer-specific chromatin interactions in established cancers.

## Methods

### Cell lines

U2OS osteosarcoma cells (HTB-96, ATCC) with a doxycycline-inducible *MYC* system were kindly provided by Elmar Wolf’s lab from Universität Würzburg, Germany. Cell lines were authenticated by STR profiling and mycoplasma testing was performed using the MycoAlert™ PLUS Mycoplasma Detection Kit (Lonza). U2OS cells were grown in DMEM supplemented with 10% *tetracycline*-free foetal bovine serum (Clontech), 100U/ml penicillin and 100µg/ml streptomycin. MYC expression was induced in U2OS cells by the addition of 1µg/ml doxycycline (Clontech) for 30h.

### Protein extraction and Western blot

Cells were lysed in RIPA Lysis and Extraction Buffer (Thermo Scientific) at 4°C for 30 minutes, with vortexing every 10 minutes. Cell lysate was centrifuged to remove cell debris. Protein concentration was measured using the Pierce BCA Protein Assay Kit (Thermo Scientific). 50 µg of protein was loaded into SDS-PAGE gel for electrophoresis. The following antibodies were used for Western blotting: anti-c-Myc (abcam, ab32072, diluted 1:1000) and anti-vinculin (Sigma, V9131, diluted 1:200) (Supplemental Table S1).

### Reverse transcription and quantitative polymerase chain reaction (RT-qPCR)

RNA was extracted from U2OS cells with or without MYC induction in triplicate, using the RNeasy Mini Kit (QIAGEN). RNA was reverse transcribed into complementary DNA using qScript cDNA Synthesis Kit (Quantabio). Quantitative PCR was performed on the QuantStudio 5 (Thermo Scientific) using the GoTaq qPCR Mastermix (Promega), with TBP as the housekeeping control. Primer sequences are listed in Supplemental Table S1.

### Circular chromosome conformation capture (4C-seq) experimental procedures

4C-seq was performed in duplicate on U2OS cells with or without *MYC* induction as previously described (Cao et al. 2017). 40 million cells were cross-linked in PBS supplemented with 1% formaldehyde at room temperature for 10 minutes, followed by quenching with 0.25M glycine for 5 minutes. Cells were washed thrice with PBS supplemented with 1mM phenylmethylsulfonyl fluoride (Sigma Aldrich). Nuclei were isolated by lysing cells in 4C Lysis Buffer (10mM Tris-HCl pH 8.0, 10mM NaCl, 5mM EDTA, 0.5% Igepal CA-630) at 4°C for 10 minutes, followed by homogenization using a dounce homogenizer (Wheaton) (50 strokes using Pestle B).

Nuclei were permeabilized at 37°C with the addition of 0.3% SDS for 1 hour, followed by 2% Triton-X 100 for 1 hour. Primary enzyme digestion with HindIII-HF (NEB) was done at 37°C for 18h, before proximity ligation in dilute conditions with T4 DNA Ligase (Thermo Scientific) at 16°C overnight. Crosslinking was reversed with the addition of 55µg/ml Proteinase K (Ambion) at 65°C for 4 hours and 37°C overnight, and treated with RNAse A at 37°C for 1 hour. DNA was purified using phenol/chloroform extraction followed by ethanol precipitation to yield the 3C library. Secondary enzyme digestion with DpnII (NEB) was done on the 3C library at 37°C overnight, followed by proximity ligation and de-crosslinking as described above. 4C libraries were generated for each viewpoint through nested inverse-PCR using Phusion DNA Polymerase (Thermo Scientific), with primers listed in Supplemental Table S1. DNA fragments between 200 and 1000bp were isolated from the 4C libraries by gel excision after running the 4C libraries on a 4-20% gradient TBE gel (Thermo Scientific). The gel slices were shredded and macerated in Tris EDTA buffer at 37°C overnight and DNA was precipitated using ethanol. The 4C libraries were multiplexed and single-end 1 x 150bp sequencing was performed on the Illumina MiSeq.

### RNA-seq experimental procedures

Total RNA was extracted from U2OS cells with or without *MYC* induction in duplicate, using the RNeasy Mini Kit (QIAGEN). Ribosomal RNA depletion and library preparation was performed using the TruSeq Stranded Total RNA LT Kit (Illumina) according to the manufacturer’s protocol. Paired-end 2 x 100bp sequencing was performed on the Illumina HiSeq 2500.

### ChIP-seq experimental procedures

H3K27ac and c-Myc ChIP-seq were performed in duplicate on U2OS cells with or without *MYC* induction. Cells were fixed in 1% formaldehyde and sonicated using the truChIP Chromatin Shearing Kit (Covaris) on the ME220 Focused-Ultrasonicator (Covaris) according to the manufacturer’s protocol. H3K27ac and c-Myc bound DNA were immunoprecipitated using anti-H3K27ac (abcam, ab4729, Lot: GR150367-1) and anti-c-Myc (Santa Cruz, sc-764, Lot: H1712), with anti-IgG (Santa Cruz, sc-2027, Lot: H2615) as the negative control (Supplemental Table S1). 3.5µg of each antibody was rotated with 15µl of Dynabeads Protein G magnetic beads (Invitrogen) at 4°C overnight and washed thrice with Beads Wash Buffer (0.1% Triton X-100 in PBS) to remove unbound antibodies. Antibody-bound beads were combined with sonicated chromatin from 5 million cells and rotated at 4°C overnight. Beads were then washed thrice with Shearing Buffer D3 (Covaris), once with High Salt Washing Buffer (50mM HEPES pH 7.5, 350mM NaCl, 1mM EDTA, 1% Triton X-100, 0.1% Sodium Deoxycholate, 0.1% SDS), once with Lithium Chloride Wash Buffer (10mM Tris pH 8.0, 250mM LiCl, 1mM EDTA, 0.5% NP-40, 0.5% Sodium Deoxycholate) and once with Tris-EDTA buffer. Chromatin was eluted from the magnetic beads in 100µl elution buffer (50mM Tris pH 8.0, 10mM EDTA, 1% SDS) supplemented with 2µl of 0.5mg/ml RNase A (QIAGEN) at 55°C for 2 hours. Chromatin was then decrosslinked with the addition of 2µl of 20mg/ml proteinase K (Ambion) at 55°C for 4 hours and 37°C overnight. For total input, chromatin from 500,000 cells were treated with RNase A and decrosslinked similarly. DNA was purified using the MinElute PCR Purification Kit (QIAGEN). ChIP-seq libraries were prepared using the ThruPLEX DNA-seq Kit (Rubicon) and sequenced paired-end 2 x 100bp on the HiSeq 2500 (Illumina).

### SIQHiC experimental procedures

SIQHiC was performed in duplicate on U2OS cells with or without *MYC* induction, with the addition of untreated mouse 3T3 cells. Human U2OS cells and mouse 3T3 cells were counted using the Countess II automated cell counter (Thermo Scientific) and fixed with 2% formaldehyde using the Arima Hi-C Kit (Arima). 1 million fixed mouse 3T3 cells were added to each sample of 4 million fixed human U2OS cells before subsequent steps of restriction enzyme digest, biotin end filling and ligation using the Arima-HiC kit (Arima Genomics) according to the manufacturer’s protocol. Libraries were prepared using the KAPA Hyper-Prep Kit (KAPA), according to the Arima-HiC kit protocol. SIQHiC libraries were sequenced paired-end 2 x 150bp on the HiSeq 4000 (Illumina).

### 4C-seq data processing

4C-seq libraries were mapped to the hg19 genome using BWA-MEM (Li 2013) version 0.7.5a-r405 using default settings and significant 4C interactions were identified using the r3Cseq pipeline (q < 0.05) (Thongjuea et al. 2013) on the CSI NGS portal (An et al. 2019) (https://csibioinfo.nus.edu.sg/). Significant 4C interactions are listed in Supplemental Table S6.

### ChIP-seq data processing

ChIP-seq libraries were mapped to the hg19 genome using BWA-MEM (Li 2013) version 0.7.5a-r405 using default settings. PCR duplicates were removed using samtools (Li et al. 2009) version 1.7 and reads falling within the ENCODE consensus blacklisted regions (Encode Project Consortium 2012) were removed using BEDTools (Quinlan and Hall 2010) version 2.26.0. ChIP-seq signals were visualized using deepTools (Ramírez et al. 2014) version 3.2.1 (bamCompare --normalizeUsing RPKM --operation subtract -bs 1). Bam files of biological replicates were merged before calling ChIP-seq peaks using MACS2 (Zhang et al. 2008) version 2.1.2. H3K27ac ChIP-seq peaks within 2kb of transcription start sites were labelled as active promoter peaks while the rest were labelled as enhancer peaks.

Superenhancers were called as described previously (Cao et al. 2017). Briefly, H3K27ac enhancer peaks within 12.5kb of each other were stitched together and ranked based on H3K27ac enrichment. The point where a line with slope 1 is tangential to the ranked H3K27ac enrichment curve was chosen as a cutoff to separate superenhancers from typical enhancers. Typical enhancers and superenhancers are listed in Supplemental Table S3.

To identify differential H3K27ac peaks, bam files from all replicates and conditions were merged and a common list of H3K27ac peaks was called using MACS2. Read counts for the common list of H3K27ac peaks were obtained using Rsubread (Liao et al. 2019). Differential H3K27ac peaks were identified using DESeq2 (Love et al. 2014) version 1.24.0 (padj<0.01 and padj<0.1 respectively).

### RNA-seq data processing

Total RNA-seq libraries were mapped to the hg19 genome and read counts for UCSC RefSeq transcripts were obtained using kallisto v0.44.0 (Bray et al. 2016). Differentially expressed transcripts were identified using sleuth v0.30.0 (Pimentel et al. 2017) (fdr<0.01, | beta| >1) and all transcript expressions are listed in Supplemental Table S4.

### SIQHiC data processing

SIQHiC libraries were analyzed using the Juicer (v1.5) pipeline (Durand et al. 2016) with some modifications to obtain hic matrix files. Since SIQHiC libraries include human and mouse DNA, paired reads were separated and mapped as single reads using BWA-MEM (Li 2013) to an artificial reference genome combining hg19 human and mm10 mouse genome sequences. Human and mouse chromosomes were appended with “H” and “M” respectively, to avoid chromosome name duplication. After mapping, reads were paired together again and split into 2 files. Paired reads both mapping to human chromosomes were placed together as Human-Human paired reads (H-H), while paired reads both mapping to mouse chromosomes were placed together as Mouse-Mouse paired reads (M-M). Ambiguous read pairs that separately mapped to different species are discarded. H-H and M-M reads were processed using Juicer separately, filtering out duplicates, intra-fragment reads and reads with MAPQ < 30 to generate .hic matrix files for each species. H-H reads of biological replicates were also merged and processed in the same way to generate .hic matrix files for downstream analyses.

The ratio between human and mouse contacts (HMR) for each sample was calculated. Since the mouse 3T3 cells in each sample come from the same population and were spiked into the human cells at the same ratio of 1:4, we expect to obtain a similar HMR ratio across all samples. Hence, the relative difference between the HMR of different samples (SIQHiC ratio) reflects the changes in global chromatin contact frequency. The SIQHiC ratio was calculated such that samples with lower HMR are normalized against samples with the highest HMR.

Hi-C contact matrices from merged biological replicates were normalized using the SIQHiC ratio by adding a custom normalization vector to the original contact matrix .hic file using the Juicer ‘addNorm’ subroutine, where the magnitude of each bin was the SIQHiC ratio raised to the power of (−0.5). In this way, the Hi-C matrices are cell-count normalized to the sample with the highest HMR to prevent scale-up extrapolation errors.

The SIQHiC normalized matrix for each chromosome was extracted from the .hic file using the Juicer ‘dump’ subroutine, combined together, and finally reassembled into a SIQHiC normalized .hic matrix file using the Juicer ‘pre’ subroutine. Since the SIQHiC normalization vector for the High *MYC* Hi-C matrix is 1, SIQHiC normalized matrix is equivalent to the non-normalized matrix. Hence, only the Low *MYC* Hi-C matrix was SIQHiC normalized. Original and SIQHiC normalized .hic matrices were balanced using the Knight-Ruiz algorithm for subsequent analyses.

Aggregate peak analyses (APA) was performed using the Juicer ‘apa’ subroutine using both non-normalized and SIQHiC normalized contact matrices using the parameters “-k KR –u –n 30”. APA heatmaps were generated using pheatmap v1.0.12 in R.

Compartments were identified using the Juicer ‘eigenvector’ and ‘pearsons’ subroutines at 100kb resolution. Since eigenvalues signs are arbitrary, we correlated eigenvectors with the number of gene promoters within each 100kb bin, such that gene rich regions were assigned positive eigenvalues.

Topologically associating domains (TADs) were identified using the Juicer Arrowhead subroutine with default parameters at 50kb and 100kb resolutions. All TADs are listed in Supplemental Table S7. Loops were called using the Juicer ‘hiccups’ subroutine with parameter “-k KR -m 1024 --ignore_sparsity”. Lost, common and gained loops were identified using pgltools (Greenwald et al. 2017) by merging and intersecting Low *MYC* loops with High *MYC* loops using a 10kb intersecting window at each loop anchor. Loop subsets are listed in Supplemental Table S5. Hi-C loops were annotated according to their overlap with c-Myc binding sites, promoters, typical enhancers and superenhancers using pgltools.

Virtual 4C tracks were extracted from the Hi-C matrices by using the Juicer ‘dump’ subroutine, dumping a 5kb window against its entire chromosome using a 5kb bin size. Virtual 4C tracks are listed in bedGraph format in Supplemental Table S6.

### Data Access

All raw and processed sequencing data generated in this study have been submitted to the NCBI Gene Expression Omnibus (GEO; https://www.ncbi.nlm.nih.gov/geo/) under accession number GSE******.

## Acknowledgements

This research is supported by the National Research Foundation (NRF) Singapore through an NRF Fellowship awarded to M.J.F (NRF-NRFF2012-054) and NTU start-up funds awarded to M.J.F. This research is supported by the RNA Biology Center at the Cancer Science Institute of Singapore, NUS, as part of funding under the Singapore Ministry of Education Academic Research Fund Tier 3 awarded to Daniel Tenen (MOE2014-T3-1-006). This research is supported by the National Research Foundation Singapore and the Singapore Ministry of Education under its Research Centres of Excellence initiative.

## Disclosure Declaration

### Ethics approval and consent to participate

Not applicable. *Consent for publication* Not applicable.

### Competing interests

M.J.F declares two patents on methodologies related to ChIA-PET. No other conflicts of interest are declared.

## References

1. An Ö, Tan K-T, Li Y, Li J, Wu C-S, Zhang B, Chen L, Yang H. 2019. CSI NGS Portal: An Online Platform for Automated NGS Data Analysis and Sharing. Preprints doi:10.20944/preprints201910.0146.v1.

2. Babu D, Fullwood MJ. 2015. 3D genome organization in health and disease: emerging opportunities in cancer translational medicine. Nucleus 6: 382–393.

3. Bahr C, von Paleske L, Uslu VV, Remeseiro S, Takayama N, Ng SW, Murison A, Langenfeld K, Petretich M, Scognamiglio R et al. 2018. A Myc enhancer cluster regulates normal and leukaemic haematopoietic stem cell hierarchies. Nature 553: 515–520.

4. Beagrie RA, Scialdone A, Schueler M, Kraemer DC, Chotalia M, Xie SQ, Barbieri M, de Santiago I, Lavitas LM, Branco MR et al. 2017. Complex multi-enhancer contacts captured by genome architecture mapping. Nature 543: 519–524.

5. Beroukhim R, Mermel CH, Porter D, Wei G, Raychaudhuri S, Donovan J, Barretina J, Boehm JS, Dobson J, Urashima M et al. 2010. The landscape of somatic copy-number alteration across human cancers. Nature 463: 899–905.

6. Boija A, Klein IA, Sabari BR, Dall’Agnese A, Coffey EL, Zamudio AV, Li CH, Shrinivas K, Manteiga JC, Hannett NM et al. 2018. Transcription Factors Activate Genes through the Phase-Separation Capacity of Their Activation Domains. Cell 175: 1842-1855.e1816.

7. Bradner JE, Hnisz D, Young RA. 2017. Transcriptional Addiction in Cancer. Cell 168: 629–643.

8. Bray NL, Pimentel H, Melsted P, Pachter L. 2016. Near-optimal probabilistic RNA-seq quantification. Nature Biotechnology 34: 525–527.

9. Busslinger GA, Stocsits RR, van der Lelij P, Axelsson E, Tedeschi A, Galjart N, Peters JM. 2017. Cohesin is positioned in mammalian genomes by transcription, CTCF and Wapl. Nature 544: 503–507.

10. Cao F, Fang Y, Tan HK, Goh Y, Choy JYH, Koh BTH, Hao Tan J, Bertin N, Ramadass A, Hunter E et al. 2017. Super-Enhancers and Broad H3K4me3 Domains Form Complex Gene Regulatory Circuits Involving Chromatin Interactions. Sci Rep 7.

11. Carter D, Chakalova L, Osborne CS, Dai YF, Fraser P. 2002. Long-range chromatin regulatory interactions in vivo. Nat Genet 32: 623–626.

12. Durand NC, Shamim MS, Machol I, Rao SSP, Huntley MH, Lander ES, Aiden EL. 2016. Juicer Provides a One-Click System for Analyzing Loop-Resolution Hi-C Experiments. Cell Systems 3: 95–98.

13. Encode Project Consortium. 2012. An integrated encyclopedia of DNA elements in the human genome. Nature 489: 57–74.

14. Gabay M, Li Y, Felsher DW. 2014. MYC Activation Is a Hallmark of Cancer Initiation and Maintenance. Cold Spring Harbor Perspectives in Medicine 4.

15. Greenwald WW, Li H, Smith EN, Benaglio P, Nariai N, Frazer KA. 2017. Pgltools: a genomic arithmetic tool suite for manipulation of Hi-C peak and other chromatin interaction data. BMC Bioinformatics 18: 207.

16. Gryder BE, Pomella S, Sayers C, Wu XS, Song Y, Chiarella AM, Bagchi S, Chou H-C, Sinniah RS, Walton A et al. 2019. Histone hyperacetylation disrupts core gene regulatory architecture in rhabdomyosarcoma. Nature Genetics 51: 1714–1722.

17. Gu B, Swigut T, Spencley A, Bauer MR, Chung M, Meyer T, Wysocka J. 2018. Transcription-coupled changes in nuclear mobility of mammalian cis-regulatory elements. Science 359: 1050–1055.

18. Heinz S, Texari L, Hayes MGB, Urbanowski M, Chang MW, Givarkes N, Rialdi A, White KM, Albrecht RA, Pache L et al. 2018. Transcription Elongation Can Affect Genome 3D Structure. Cell 174: 1522-1536.e1522.

19. Herranz D, Ambesi-Impiombato A, Palomero T, Schnell SA, Belver L, Wendorff AA, Xu L, Castillo-Martin M, Llobet-Navás D, Cordon-Cardo C et al. 2014. A NOTCH1-driven MYC enhancer promotes T cell development, transformation and acute lymphoblastic leukemia. Nature Medicine 20: 1130–1137.

20. Hnisz D, Abraham Brian J, Lee Tong I, Lau A, Saint-André V, Sigova Alla A, Hoke Heather A, Young Richard A. 2013. Super-Enhancers in the Control of Cell Identity and Disease. Cell 155: 934–947.

21. Isoda T, Moore AJ, He Z, Chandra V, Aida M, Denholtz M, Piet van Hamburg J, Fisch KM, Chang AN, Fahl SP et al. 2017. Non-coding Transcription Instructs Chromatin Folding and Compartmentalization to Dictate Enhancer-Promoter Communication and T Cell Fate. Cell 171: 103-119.e118.

22. Kieffer-Kwon K-R, Nimura K, Rao SSP, Xu J, Jung S, Pekowska A, Dose M, Stevens E, Mathe E, Dong P et al. 2017. Myc Regulates Chromatin Decompaction and Nuclear Architecture during B Cell Activation. Molecular Cell 67: 566-578.e510.

23. Kress TR, Sabo A, Amati B. 2015. MYC: connecting selective transcriptional control to global RNA production. Nat Rev Cancer 15: 593–607.

24. Li H. 2013. Aligning sequence reads, clone sequences and assembly contigs with BWA-MEM. eprint arXiv:13033997: 1303.3997.

25. Li H, Handsaker B, Wysoker A, Fennell T, Ruan J, Homer N, Marth G, Abecasis G, Durbin R. 2009. The Sequence Alignment/Map format and SAMtools. Bioinformatics (Oxford, England) 25: 2078–2079.

26. Liao Y, Smyth GK, Shi W. 2019. The R package Rsubread is easier, faster, cheaper and better for alignment and quantification of RNA sequencing reads. Nucleic Acids Research 47: e47–e47.

27. Lin CY, Loven J, Rahl PB, Paranal RM, Burge CB, Bradner JE, Lee TI, Young RA. 2012. Transcriptional amplification in tumor cells with elevated c-Myc. Cell 151: 56–67.

28. Lorenzin F, Benary U, Baluapuri A, Walz S, Jung LA, von Eyss B, Kisker C, Wolf J, Eilers M, Wolf E. 2016. Different promoter affinities account for specificity in MYC-dependent gene regulation. eLife 5: e15161.

29. Love MI, Huber W, Anders S. 2014. Moderated estimation of fold change and dispersion for RNA-seq data with DESeq2. Genome Biology 15: 550.

30. Loven J, Hoke HA, Lin CY, Lau A, Orlando DA, Vakoc CR, Bradner JE, Lee TI, Young RA. 2013. Selective inhibition of tumor oncogenes by disruption of super-enhancers. Cell 153: 320–334.

31. Loven J, Orlando DA, Sigova AA, Lin CY, Rahl PB, Burge CB, Levens DL, Lee TI, Young RA. 2012. Revisiting global gene expression analysis. Cell 151: 476–482.

32. Lovén J, Orlando David A, Sigova Alla A, Lin Charles Y, Rahl Peter B, Burge Christopher B, Levens David L, Lee Tong I, Young Richard A. 2012. Revisiting Global Gene Expression Analysis. Cell 151: 476–482.

33. Lun ATL, Smyth GK. 2015. diffHic: a Bioconductor package to detect differential genomic interactions in Hi-C data. BMC Bioinformatics 16: 258.

34. McKeown MR, Bradner JE. 2014. Therapeutic strategies to inhibit MYC. Cold Spring Harb Perspect Med 4.

35. Nair SK, Burley SK. 2003. X-Ray Structures of Myc-Max and Mad-Max Recognizing DNA: Molecular Bases of Regulation by Proto-Oncogenic Transcription Factors. Cell 112: 193–205.

36. Nie Z, Hu G, Wei G, Cui K, Yamane A, Resch W, Wang R, Green DR, Tessarollo L, Casellas R et al. 2012. c-Myc is a universal amplifier of expressed genes in lymphocytes and embryonic stem cells. Cell 151: 68–79.

37. Ooi WF, Xing M, Xu C, Yao X, Ramlee MK, Lim MC, Cao F, Lim K, Babu D, Poon L-F et al. 2016. Epigenomic profiling of primary gastric adenocarcinoma reveals super-enhancer heterogeneity. Nature Communications 7: 12983.

38. Orlando David A, Chen Mei W, Brown Victoria E, Solanki S, Choi Yoon J, Olson Eric R, Fritz Christian C, Bradner James E, Guenther Matthew G. 2014. Quantitative ChIP-Seq Normalization Reveals Global Modulation of the Epigenome. Cell Reports 9: 1163–1170.

39. Pimentel H, Bray NL, Puente S, Melsted P, Pachter L. 2017. Differential analysis of RNA-seq incorporating quantification uncertainty. Nature Methods 14: 687–690.

40. Plank Jennifer L, Dean A. 2014. Enhancer Function: Mechanistic and Genome-Wide Insights Come Together. Molecular Cell 55: 5–14.

41. Quinlan AR, Hall IM. 2010. BEDTools: a flexible suite of utilities for comparing genomic features. Bioinformatics (Oxford, England) 26: 841–842.

42. Raisner R, Bainer R, Haverty PM, Benedetti KL, Gascoigne KE. 2020. Super-enhancer acquisition drives oncogene expression in triple negative breast cancer. PLOS ONE 15: e0235343.

43. Ramírez F, Dündar F, Diehl S, Grüning BA, Manke T. 2014. deepTools: a flexible platform for exploring deep-sequencing data. Nucleic Acids Research 42: W187–W191.

44. Sabari BR, Dall Agnese A, Boija A, Klein IA, Coffey EL, Shrinivas K, Abraham BJ, Hannett NM, Zamudio AV et al. 2018. Coactivator condensation at super-enhancers links phase separation and gene control. Science 361: 379-+.

45. Sabo A, Kress TR, Pelizzola M, de Pretis S, Gorski MM, Tesi A, Morelli MJ, Bora P, Doni M, Verrecchia A et al. 2014. Selective transcriptional regulation by Myc in cellular growth control and lymphomagenesis. Nature 511: 488–492.

46. Sati S, Cavalli G. 2017. Chromosome conformation capture technologies and their impact in understanding genome function. Chromosoma 126: 33–44.

47. Schaub FX, Dhankani V, Berger AC, Trivedi M, Richardson AB, Shaw R, Zhao W, Zhang X, Ventura A, Liu Y et al. 2018. Pan-cancer Alterations of the MYC Oncogene and Its Proximal Network across the Cancer Genome Atlas. Cell Systems 6: 282-300.e282.

48. Schmitt AD, Hu M, Jung I, Xu Z, Qiu Y, Tan CL, Li Y, Lin S, Lin Y, Barr CL et al. 2016. A Compendium of Chromatin Contact Maps Reveals Spatially Active Regions in the Human Genome. Cell Rep 17: 2042–2059.

49. Shi J, Whyte WA, Zepeda-Mendoza CJ, Milazzo JP, Shen C, Roe J-S, Minder JL, Mercan F, Wang E, Eckersley-Maslin MA et al. 2013. Role of SWI/SNF in acute leukemia maintenance and enhancer-mediated Myc regulation. Genes & Development 27: 2648–2662.

50. Shrivastava A, Yu J, Artandi S, Calame K. 1996. YY1 and c-Myc associate in vivo in a manner that depends on c-Myc levels. Proceedings of the National Academy of Sciences 93: 10638.

51. Stansfield JC, Cresswell KG, Vladimirov VI, Dozmorov MG. 2018. HiCcompare: an R-package for joint normalization and comparison of HI-C datasets. BMC Bioinformatics 19: 279.

52. Thongjuea S, Stadhouders R, Grosveld FG, Soler E, Lenhard B. 2013. r3Cseq: an R/Bioconductor package for the discovery of long-range genomic interactions from chromosome conformation capture and next-generation sequencing data. Nucleic Acids Research 41: e132–e132.

53. Tsang FH, Law CT, Tang TC, Cheng CL, Chin DW, Tam WV, Wei L, Wong CC, Ng IO, Wong CM. 2019. Aberrant Super-Enhancer Landscape in Human Hepatocellular Carcinoma. Hepatology (Baltimore, Md) 69: 2502–2517.

54. Vella P, Barozzi I, Cuomo A, Bonaldi T, Pasini D. 2012. Yin Yang 1 extends the Myc-related transcription factors network in embryonic stem cells. Nucleic Acids Research 40: 3403–3418.

55. Walz S, Lorenzin F, Morton J, Wiese KE, von Eyss B, Herold S, Rycak L, Dumay-Odelot H, Karim S, Bartkuhn M et al. 2014. Activation and repression by oncogenic MYC shape tumour-specific gene expression profiles. Nature 511: 483–487.

56. Weintraub AS, Li CH, Zamudio AV, Sigova AA, Hannett NM, Day DS, Abraham BJ, Cohen MA, Nabet B, Buckley DL et al. 2017. YY1 Is a Structural Regulator of Enhancer-Promoter Loops. Cell 171: 1573-1588.e1528.

57. Whyte Warren A, Orlando David A, Hnisz D, Abraham Brian J, Lin Charles Y, Kagey Michael H, Rahl Peter B, Lee Tong I, Young Richard A. 2013. Master Transcription Factors and Mediator Establish Super-Enhancers at Key Cell Identity Genes. Cell 153: 307–319.

58. Zeid R, Lawlor MA, Poon E, Reyes JM, Fulciniti M, Lopez MA, Scott TG, Nabet B, Erb MA, Winter GE et al. 2018. Enhancer invasion shapes MYCN-dependent transcriptional amplification in neuroblastoma. Nature Genetics 50: 515–523.

59. Zhang X, Choi PS, Francis JM, Imielinski M, Watanabe H, Cherniack AD, Meyerson M. 2016. Identification of focally amplified lineage-specific super-enhancers in human epithelial cancers. Nature Genetics 48: 176–182.

60. Zhang Y, Liu T, Meyer CA, Eeckhoute J, Johnson DS, Bernstein BE, Nusbaum C, Myers RM, Brown M, Li W et al. 2008. Model-based Analysis of ChIP-Seq (MACS). Genome Biology 9: R137.

